# *Nepeta cataria* L. (catnip) can serve as a chassis for the engineering of secondary metabolic pathways

**DOI:** 10.1101/2023.10.11.561176

**Authors:** Marcus Geissler, Christoph Neubauer, Yuriy V. Sheludko, Adrian Brückner, Heribert Warzecha

## Abstract

**Objective:** Evaluation of *Nepeta cataria* as a host with specific endogenous metabolite background for transient expression and metabolic engineering of secondary biosynthetic sequences.

**Results:** The reporter gene *GFP::lic*BM3 as well as three biosynthetic genes leading to the formation of the cannabinoid precursor olivetolic acid were adopted to the modular cloning standard GoldenBraid, transiently expressed in two chemotypes of *N. cataria* and compared to *Nicotiana benthamiana*. To estimate the expression efficiency in both hosts, quantification of the reporter activity was carried out with a sensitive and specific lichenase assay. While *N. benthamiana* exhibited lichenase activity of 676 ± 94 μmol g^-1^ s^-1^ (Gerasimenko et al. 2019), *N. cataria* cultivar ‘1000’, and the cultivar ‘Citriodora’ showed an activity of 37 ± 8 μmol g^-1^ s^-1^ and 18 ± 4 μmol g^-1^ s^-1^, respectively. Further, combinatorial expression of genes involved in cannabinoid biosynthetic pathway *acylactivating enzyme 1* (*aae1*), *olivetol synthase* (*ols*) and *olivetolic acid cyclase* (*oac*) in *N. cataria* cv. resulted presumably in the *in vivo* production of olivetolic acid glycosides.

**Conclusion:** *Nepeta cataria* is amenable to *Agrobacterium*-mediated transient expression and could serve as a novel chassis for the engineering of secondary metabolic pathways and transient evaluation of heterologous genes.

## Introduction

Engineering of metabolic pathways in alternative hosts holds a promising strategy to improve the production of valuable secondary metabolites or to generate new-to-nature compounds (Luo et al. 2015). *Nicotiana benthamiana* Domin has become both a model system and working horse for transient gene expression toward recombinant protein production as well as a platform for enzymatic pathway assembly and metabolite production (Bally et al. 2018). This fact also makes *N. benthamiana* an ideal exploration tool to develop novel metabolic engineering strategies, evaluate enzymatic cascades, and enable the fast and reliable testing of genetic constructs prior to a time-consuming stable transformation.

Although *N. benthamiana* has successfully been deployed to host complex metabolic routes and produce natural products of interest (Grzech et al. 2023) it has some limitations. Unintended glycosylation of pathway intermediates or end products (Glück et al. 2020) or proteolysis of recombinant proteins (Grosse-Holz et al. 2017) has been described, which are an obstacle for high yields of any given product demanding complex and intensive engineering of the host genome (Dudley et al. 2022). Another factor that might limit the capacity of *N. benthamiana* is precursor supply, necessary for fueling the pipeline leading to secondary metabolites. The assembly of these specialized compounds rely on the supply of building blocks deviated from primary metabolism, and this is especially true for natural products containing terpenoid moieties within their structures, like monoterpenoid indole alkaloids (Geissler et al. 2016) or terpenophenolics (Schachtsiek et al. 2017). Hence, a strong endogenous supply of precursors like isopentenyl diphosphate (IPP) or geranyl diphosphate (GPP) in the heterologous host plant might be an advantage for the formation of novel metabolites. Moreover, sequestration and storage of newly produced metabolites seems to be a crucial aspect of high-level metabolite formation in host organisms. Especially glandular trichomes fulfill the role of specialized organs optimized to produce and store extreme amounts of natural products incompatible with the aqueous environment within a cell (Huchelmann et al. 2017).

Considering the reasons mentioned above, we identified *Nepeta cataria* L. (catnip) as a potential host plant for metabolic engineering efforts because of its chemotypes, rich in terpenoid constituents (nepetalactones and iridoids or citral derivatives (Said-Al Ahl et al. 2018)), and well-developed trichome system for the storage and secretion of the volatile components of essential oils. Hence, *N. cataria* could serve as a host for enzymatic routes relying on the terpenoid precursors. An important example of such a route is cannabinoid metabolic pathway, involving the integration of polyketides and terpene metabolites derived from the methylerythritol 4-phosphate pathway (Fellermeier et al. 2001; Gülk and Möller 2020; Supplementary Figure 1

**Fig. 1.**
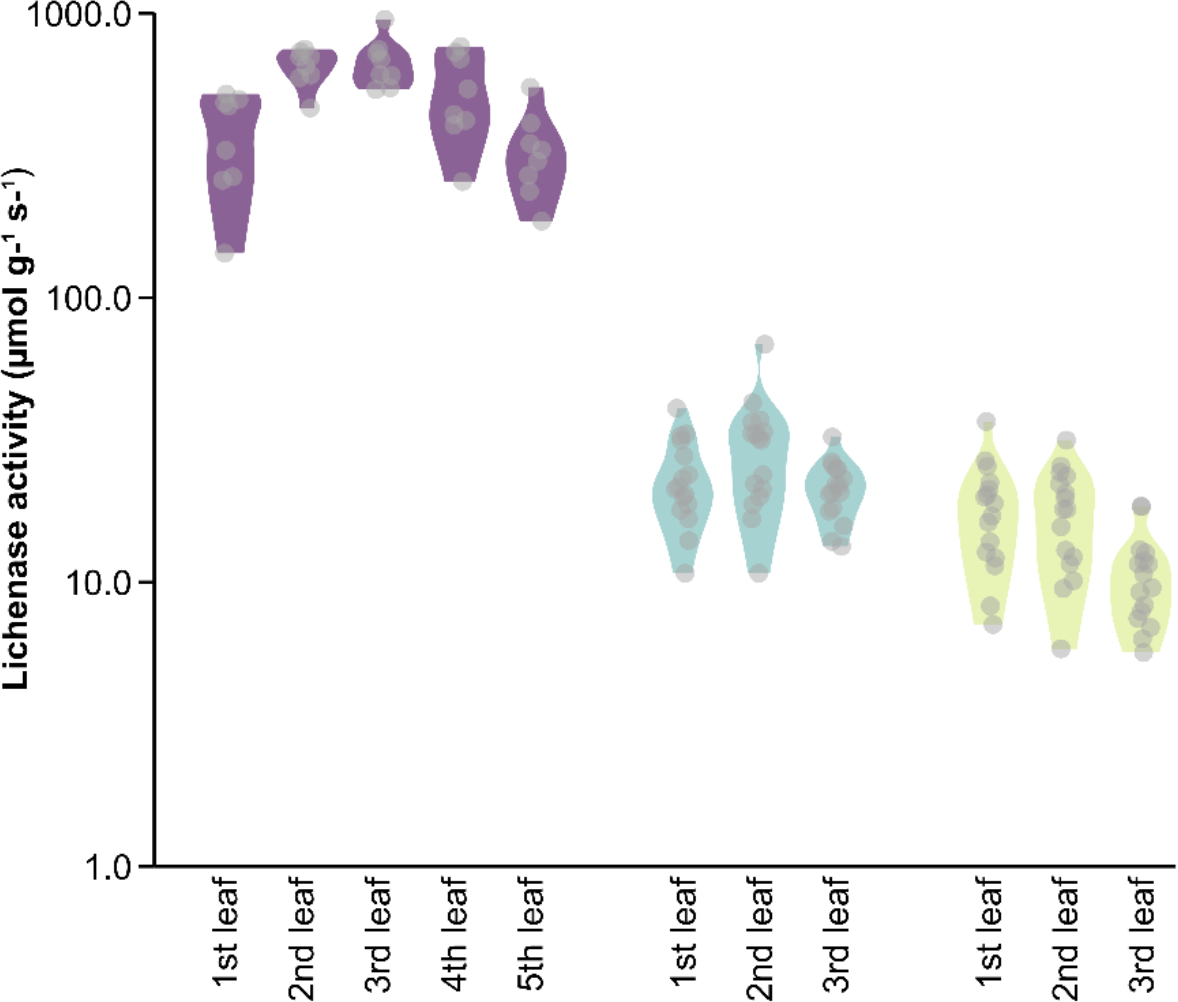
Lichenase activity in *N. benthamiana* (Gerasimenko et al., 2019; purple) and *N. cataria* leaves (cultivar ‘1000’ in teal and cultivar ‘Citriodora’ in lime) of different ages (1st-, the youngest leaf). In case of catnip plants, the two upper, the two middle and the two lower leaves were summarized as the first, second and third leaf, respectively, since they are emerging from the same node. Jittered dots represent the measured values, while the violines depict the data distribution for each leaf type.

Here we evaluate two chemotypes of *N. cataria* for their suitability for transient expression of heterologous genes, involved in a polyketide branch of cannabinoid biosynthesis, and accumulation of cannabinoid intermediates.

## Materials and Methods Chemicals

Nerol, β-citronellol, geraniol and citral, used as authentic standards in GC–MS analysis, were procured from Sigma-Aldrich (St. Louis, MO, USA). Olivetolic acid (OA; Santa Cruz Biotechnology, Heidelberg, Germany) served as an authentic standard as well as a substrate in supplementation experiments. Hexanoic acid, used as precursor supply, was purchased from Carl Roth (Karlsruhe, Germany).

### Bacterial strains and growth conditions

*Escherichia coli* TOP10 cells (Thermo Fisher Scientific, Waltham, USA), used for cloning, and *Agrobacterium tumefaciens* strain EHA105 and GV3101 (ICON Genetics, Halle, Germany), applied in transient expression experiments, were maintained as described in Fräbel et al. (2016).

### Cloning of DNA

To generate the genetic constructs used for transient transformation of *N. cataria* and *N. benthamiana* plants, restriction-ligation reactions were set up as reported by (Sarrion-Perdigones et al. 2013). The domestication of *aae1* (GenBank AFD33345.1), *ols* (GenBank AB164375.1) and *oac* (GenBank AFN42527.1), as well as the generation of *A. tumefaciens* EHA105 and GV3101 for transformation of plants is described in more detail in the supporting information. The constructs were assembled the way as described by Fräbel et al. (2016) utilizing the cauliflower mosaic virus 35S promoter (P35S) and the nopaline synthase terminator (TNos) for regulation of gene expression.

As a bifunctional reporter, we utilized an optimized version of *lic*BM3 gene from *Clostridium thermocellum* (*Ruminiclostridium thermocellum*), coding for thermostable lichenase, in translational fusion with a synthetic GFP gene (NCBI acc. no KX458181; Gerasymenko et al. 2017).

### Transient transformation of plants

Transient transformation of greenhouse-grown *N. cataria* and *N. benthamiana* plants was performed as previously described (Geissler et al. 2018). For reconstruction of the OA biosynthetic pathway, plants were additionally injected with 4 mM of hexanoic acid (solved in infiltration buffer (10 mM MES, 10 mM MgSO_4_) four days after initial infiltration with agrobacteria harboring *aae1, ols* and *oac* and harvested after 24 h of incubation.

### Determination of lichenase activity

Total soluble proteins (TSP) were obtained as previously described (Geissler et al. 2018) using approximately 80 mg of plant material. For the determination of lichenase activity, assays were performed as described by (Gerasimenko et al., 2019).

### GC*–*MS analysis of volatile compounds

For the analysis of volatiles accumulated in *N. cataria* plant material, the compounds were extracted with dichloromethane (DCM, Carl Roth, Karlsruhe, Germany) and the obtained samples measured by GC–MS using a DB-5MS column (30 m, 0.25 mm ID, 0.25 μm film thickness; Phenomenex, Aschaffenburg, Germany). Detailed parameters of sample preparation and GC–MS analysis are specified in the supplementary data.

### Extraction of metabolites and HPLC–MS analysis of cannabinoid precursors

150 mg of frozen plant material was homogenized by sonication in 200 μl of methanol/water (80:20, by vol.) for 30 min. The extracts were cleared by centrifugation at 17,000 ×*g* for 10 min and 4 °C and the supernatant subjected to HPLC−MS analysis using a Poroshell 120SB-C18 (3.0×150 mm, 2.7 μm; Agilent, Santa Clara, CA, USA) column, as previously described (Geissler et al. 2018). OA and OA glucosides were detected in negative selected ion monitoring (SIM) with selected m/z of 223.2 and 385.2, respectively.

## Results and discussion

### *Agrobacterium*-mediated transient transformation of *N. cataria* plants

In the first step, we wanted to evaluate the amenability of catnip to the infiltration with an *Agrobacterium* suspension. Secondly, heterologous gene expression needs to be verified by fluorescence microscopy and by enzymatic assay of lichenase activity. The upper six leaves of four-week-old plants were grouped in pairs according to the nodes from which they emerged. Plants were infiltrated either with *A. tumefaciens* (strain GV3101) or with the same strain harbouring the GFP:LicBM3 expression construct, providing a fusion of the *gfp* gene for fluorescence readout and the *licBM3* gene encoding the enzyme lichenase whose activity can be quantified after successful expression (Supplementary Figure 3a). Additionally, a transcriptional unit harbouring the p19 protein to avoid post-transcriptional gene silencing was included in the vector (Supplementary Figure 2). After seven days of incubation, all plants displayed barely any necrosis compared to non-infiltrated leaves (Supplementary Figure 3b and c). Under UV light, an easily visible fluorescence could be observed in plants transformed with GV3101 carrying the GFP:LicBM3 expression construct (Supplementary Figure 4). Since the transient transformation of *N. cataria* plants visually proved possible, a quantitative analysis of transformation efficiency was performed next. Lichenase activity was measurable in all three leaf pairs of both chemovars with 37 ± 8 μmol g^-1^ s^-1^ for the second leaf pair of *N. cataria* ‘1000’, and 18 ± 4 μmol g^-1^ s^-1^ activity in the first, and thus youngest, leaf pair of *N. cataria* ‘Citriodora’ (Figure 1). Activity differed between leaves, but the difference was, however, not significant. For comparison, we took the data of *N. benthamiana*, four-week-old plants transformed with the same construct (Gerasymenko et al. 2019), where lichenase activity is most pronounced in the second to third upper leaves (650 ± 65 and 676 ± 94 μmol g^-1^ s^-1^, respectively, Figure 1).

Although the expression level of the reporter protein in *N. benthamiana* was considerably higher compared to *N. cataria*, the general infiltration approach proved viable for both tested *N. cataria* varieties providing a detectable yield of the reporter protein. It cannot be excluded that other developmental stages of the plants could be more optimal concerning the expression levels since plant senescence alters gene expression and the endogenous phytohormone balance, which may influence susceptibility to viral/bacterial infection (Buchanan-Wollaston et al. 2005). Given the fact that older leaves were less amenable to the infiltration procedure and that the initial results confirmed the potential of catnip for rapid testing of genetic sequences for metabolite production, we continued with the established protocol.

### Characterization of chemotypes within the different *N. cataria* cultivars

The *Nepeta* genus has been the subject of extensive research and studies on their phytochemical variations led to the definition of two main catnip chemotypes. One, with nepetalactones as the dominant compounds, and another with citral derivatives as major constituents (Said-Al Ahl et al. 2018). To evaluate the effect of the endogenous metabolic background on any heterologous pathway being introduced it was necessary to characterize the two utilized cultivars regarding their chemical profiles. Therefore, volatile compounds were extracted from the two studied *N. cataria* cultivars and analyzed by GC–MS. The investigation revealed two distinct chemotypes. The cultivar ‘Citriodora’ consisted exclusively of chemotype 1, containing citral derivatives like citronellol, neral, geraniol and geranial, while *N. cataria* cultivar ‘1000’ consisted exclusively of chemotype 2 containing diastereomers of nepetalactone in different proportions (Figure 2).

**Fig. 2.**
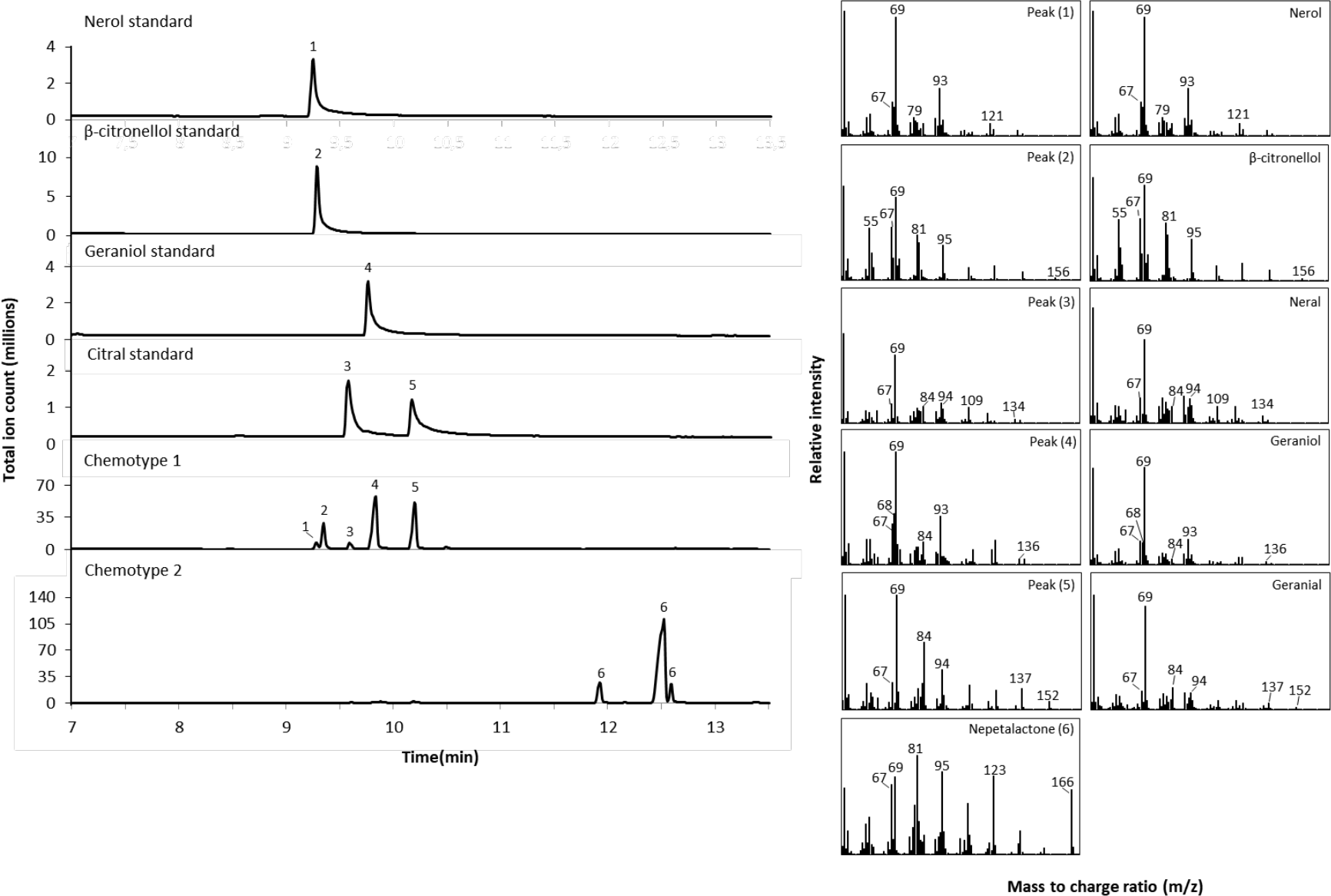
GC−MS analysis of volatile compounds produced in two different *N. cataria* wildtype (WT) cultivars, consisting mainly of either citral derivatives (chemotype 1) or nepetalactone (chemotypes 2). 1, nerol; 2, β-citronellol; 3, neral; 4, geraniol; 5, geranial; 6, nepetalactone. The identities of metabolites 1-5 were confirmed by comparison of their MS spectra with authentic standards. Metabolite 6 was verified by comparison of its MS spectrum with reference outputs deposited in the National Institute of Standards and Technology (NIST) database and represents nepetalactone (mixture of diastereomers).

### Production of olivetolic acid (OA) metabolites in transiently transformed *N. cataria* plants

For the introduction of a new pathway, we chose the enzymatic route derived from *Cannabis sativa* leading to the formation of phytocannabinoids. Although this pathway starts from an unusual precursor, hexanoic acid, it is well suited to probe our assumption of *N. cataria* capable of serving as a host for metabolic engineering efforts. As terpenophenolics, cannabinoids combine two biosynthetic routes, merging polyketides with isoprenoids (Schachtsiek et al. 2017). Here, we started with the first three enzymatic steps leading to the formation of olivetolic acid (OA), a key polyketide intermediate in the cannabinoid biosynthetic pathway.

As a prerequisite of the metabolic engineering efforts, we determined whether OA has toxic effects in catnip, as it has been shown for cannabigerolic acid (CBGA) and Δ^9^-tetrahydrocannibinolic acid (THCA) fed to *C. sativa* cell suspension cultures (Sirikantaramas et al. 2005). Within the same experimental setting, the suitability of the extraction method for isolation of OA could be evaluated. Therefore, *N. cataria* plants were injected with OA and incubated for 24 h. After extraction and analysis by HPLC-MS, not only OA could be detected, but also two other metabolites which eluted approximately 2 min earlier, exhibiting an *m/z* [M-H]^-^ 385.2 (Figure 3), suggesting that the extracted substances are the glycosylated forms of OA. This was also described for *N. bethamiana* transiently expressing the pathway genes to produce OA and whereupon the OA formed was glucosylated either at the hydroxy group at C-2 or at C-4 *in planta* (Gülck et al. 2020; Figure 3).

**Fig. 3.**
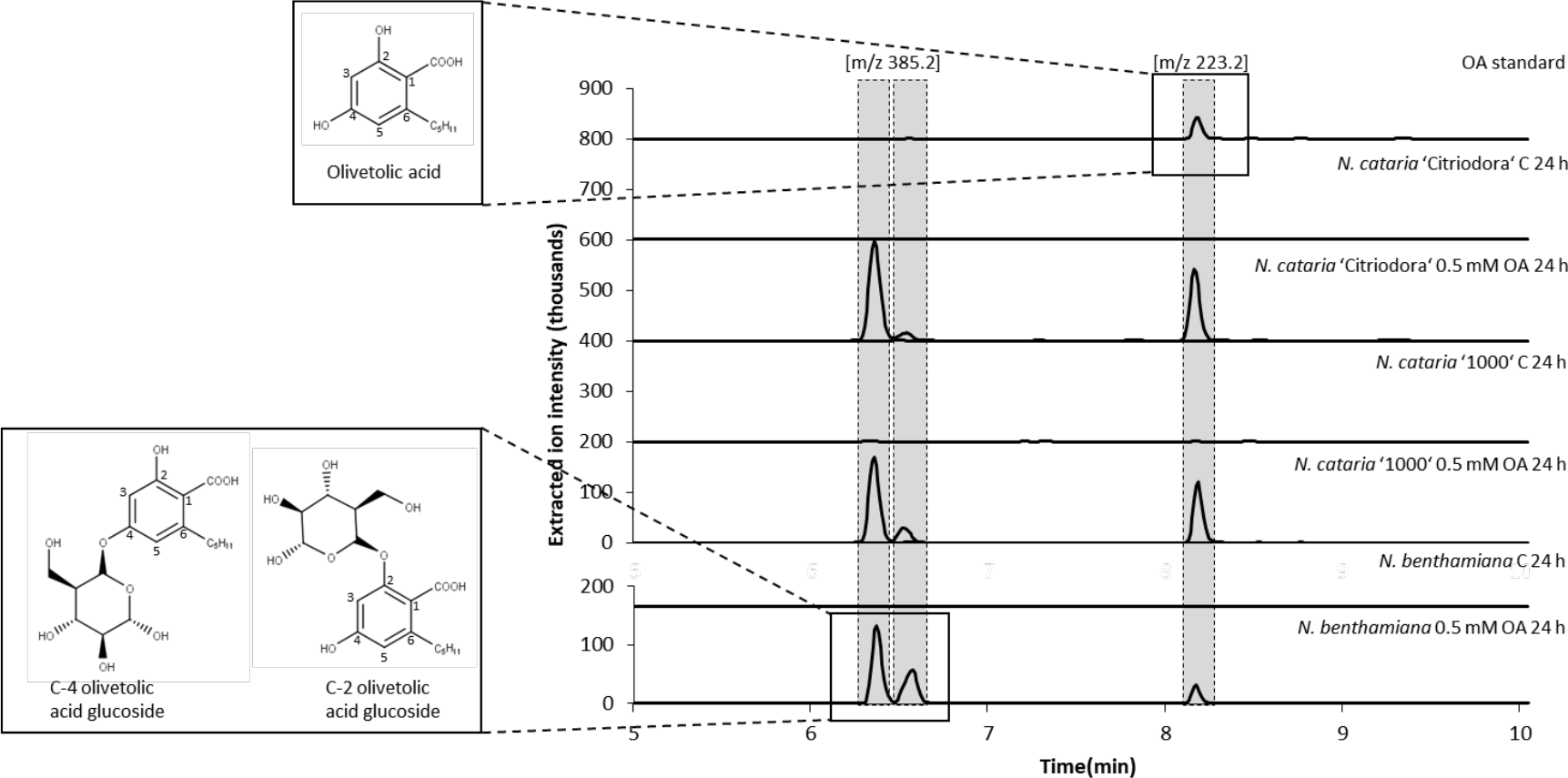
HPLC−MS analysis of *N. cataria* cultivar ‘Citriodora’ and ‘1000’ as well as *N. benthamiana* plant extracts obtained from leaves infiltrated with olivetolic acid (OA). As a negative control (C) wildtype plants were solely infiltrated with infiltration buffer. In addition to OA, two additional metabolites with an m/z of 385.2 were identified. In comparison with the results obtained by Gülck et al. (2020) with OA infiltrated into *Nicotiana benthamiana* plants, the metabolites eluting might represent C-4 OA glucoside and C-2 OA glucoside.

Next, metabolic engineering of OA biosynthesis had to be achieved by introduction of the pathway genes into the plant host. Therefore, genetic constructs were assembled for the accumulation of the desired enzymes (AAE1, OAC and OLS) and p19 suppressor of gene silencing in the cytosol (Supplementary Figure 2; construct TOA1 and TOA2). For transient transformation of *N. cataria* plants, *Agrobacterium* suspensions harboring the constructs TOA1 and TOA2 were co-infiltrated, supplemented with 4 mM hexanoic acid four days post infiltration and harvested after another 24 h of incubation. Transiently transformed plants solely infiltrated with the infiltration buffer were used as a negative control. The extracted metabolites were administered to HPLC−MS and analyzed in terms of the production of OA and its glycosylated forms. Expression of the transgenes resulted in the detection of OA glycoside when supplemented with hexanoic acid (Figure 4), indicating efficient assembling of the biosynthetic sequence in *N. cataria*. Detailed structure elucidation of the formed glycosides by NMR spectroscopy might be possible after preparative isolation of the products and is beyond the scope of this paper.

In conclusion, using the polyketide branch of cannabinoid biosynthesis as an example, we have shown for the first time that catnip can be used as a chassis for the assembling of heterologous metabolic pathway and production of PNPs, representing an alternative to the model plant *N. benthamiana*.

**Fig. 4.**
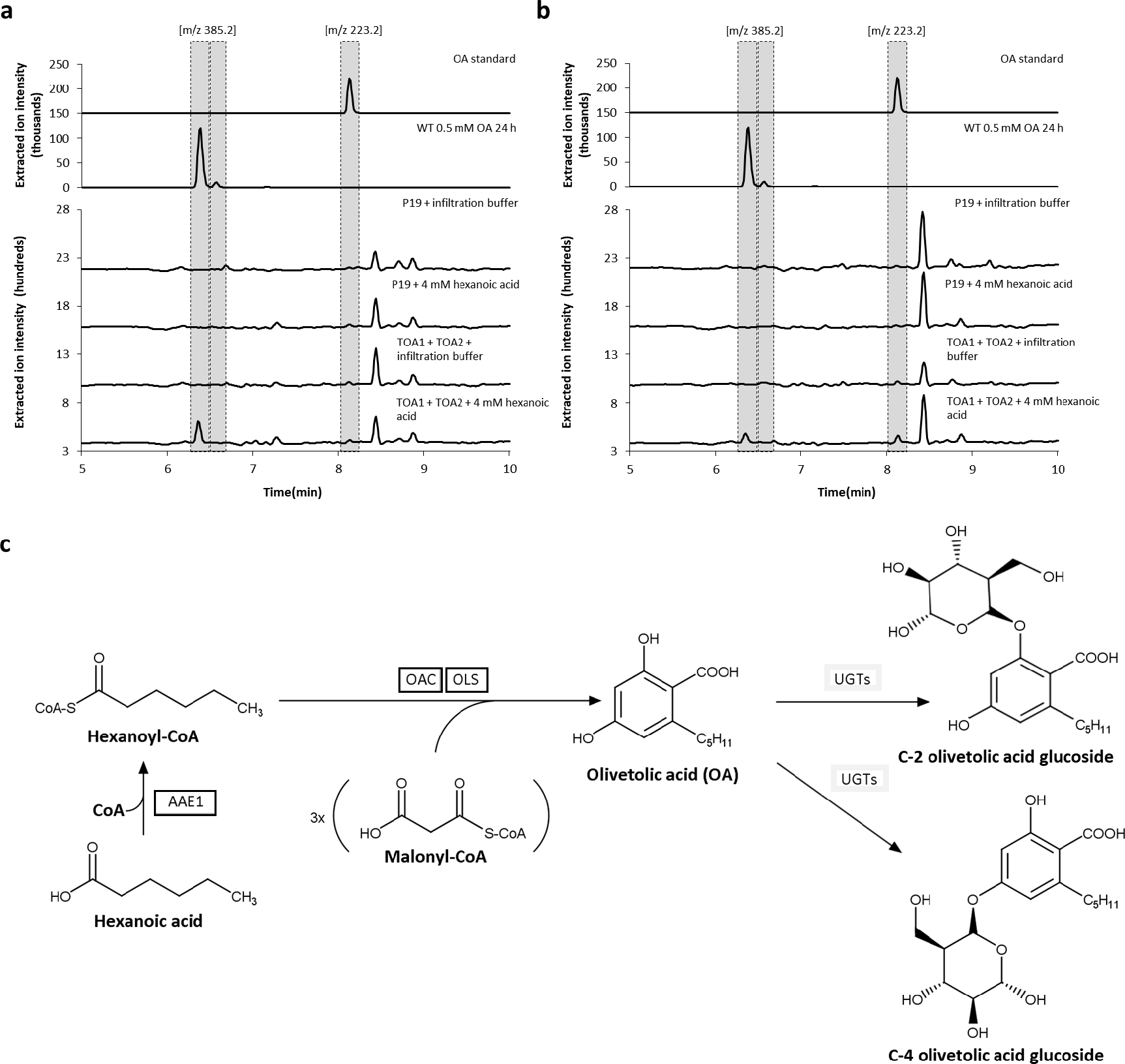
HPLC−MS analysis of *N. cataria* cultivar ‘1000’ (a) and ‘Citriodora’ (b) plant extracts obtained from leaves expressing the pathway genes required for the production of olivetolic acid (OA) and additional supplementation of 4 mM hexanoic acid. *N. cataria* wildtype plants infiltrated with OA served as a positive control. Plants infiltrated with infiltration buffer as well as plants injected with *Agrobacterium* harboring p19 and supplemented with hexanoic acid were used as negative controls. (c) Schematic depiction of the biosynthetic pathway leading to glucosylated olivetolic acid. AAE1, acyl-activating enzyme 1; OLS, olivetol synthase; OAC, olivetolic acid cyclase; UGTs, endogenous UDP-glycosyltransferases. Heterologously produced enzymes are boxed.

## Supporting information

Supplementary Fig. 1 Schematic depiction of the cannabinoid biosynthesis pathway in Cannabis sativa. MEP, 2-C-methyl-D-erythritol-4-phosphate pathway; AAE1, acyl activating enzyme 1; OLS, olivetol synthase; OAC, olivetolic acid cyclase; CBGAS, cannabigerolic acid synthase; THCAS, Δ9-tetrahydrocannabinolic acid synthase; CBDAS, cannabidiolic acid synthase; CBCAS, cannabichromenic acid synthase.

Supplementary Fig. 2 - Schematic representation of the generated constructs within the GoldenBraid grammar (Sarrion-Perdigones et al. 2013). The capital letters show the four−nucleotide overhangs ensuring correct final orientation within the transcriptional unit (TU), while the numbers above the scheme represent standard GoldenBraid classes within the TU structure. P35S ATG, cauliflower mosaic virus (CaMV) 35S promoter with an integrated start codon ensuring cytosolic localization; TNos, nopaline synthase terminator; 8×his:TNos, nopaline synthase terminator comprising an 8×his-tag; *aae*1, acyl-activating enzyme 1; *ols*, olivetol synthase; *oac*, olivetolic acid cyclase; *gfp:lic*BM3, lichenase in translational fusion to a synthetic GFP. Abbreviation of each construct is listed on the right. Boxes are not drawn to scale.

Supplementary Fig. 3 (a) Infiltration scheme of the upper six leaves (1a, 1b, 2a, 2b, 3a, 3b) of four-week-old *N. cataria* plants. (b) The infiltrated leaves showed very little necrosis at seven dpi compared to non-infiltrated leaves (c).

Supplementary Fig. 4 Fluorescence microscopy of *N. cataria* variety ‘Citriodora’ and ‘1000’ and *N. benthamiana* plants expressing *gfp:licBM3+P19*. Infiltration of plants with untransformed *Agrobacterium* served as a negative control (C).

## Notes

### Competing Interest Statement

The authors have declared no competing interest.

