## Supplements for "*Nepeta cataria* L. (catnip) can serve as a chassis for the engineering of secondary metabolic pathways"

### Captions

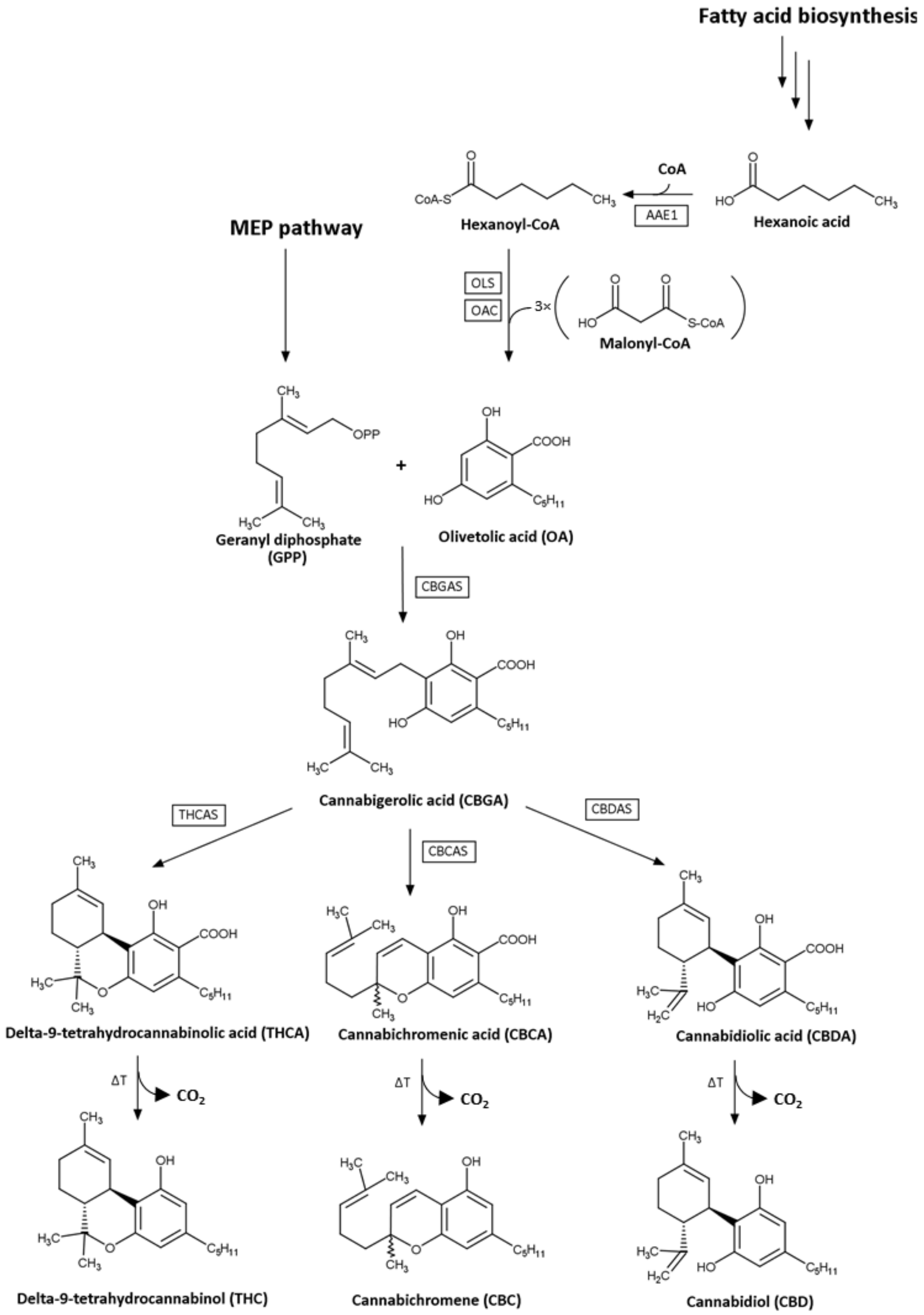


**Supplementary Fig. 1** Schematic depiction of the cannabinoid biosynthesis pathway in Cannabis sativa. MEP, 2-C-methyl-D-erythritol-4-phosphate pathway; AAE1, acyl activating enzyme 1; OLS, olivetol synthase; OAC, olivetolic acid cyclase; CBGAS, cannabigerolic acid synthase; THCAS, Δ9-tetrahydrocannabinolic acid synthase; CBDAS, cannabidiolic acid synthase; CBCAS, cannabichromenic acid synthase.

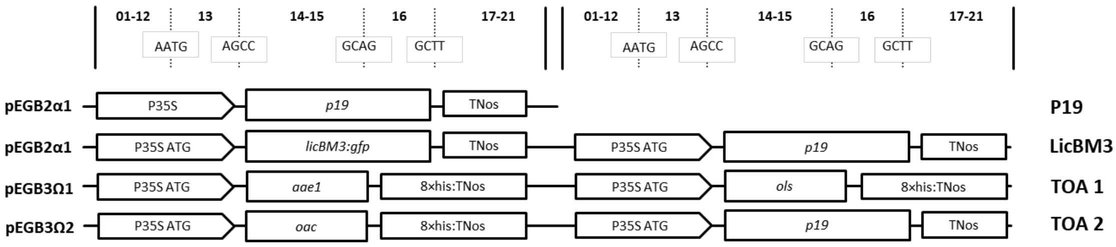


**Supplementary Fig. 2** Schematic representation of the generated constructs within the GoldenBraid grammar (Sarrion-Perdigones et al. 2013). The capital letters show the four−nucleotide overhangs ensuring correct final orientation within the transcriptional unit (TU), while the numbers above the scheme represent standard GoldenBraid classes within the TU structure. P35S ATG, cauliflower mosaic virus (CaMV) 35S promoter with an integrated start codon ensuring cytosolic localization; TNos, nopaline synthase terminator; 8×his:TNos, nopaline synthase terminator comprising an 8×his-tag; *aae*1, acyl-activating enzyme 1; *ols*, olivetol synthase; *oac*, olivetolic acid cyclase; *gfp:lic*BM3, lichenase in translational fusion to a synthetic GFP. Abbreviation of each construct is listed on the right. Boxes are not drawn to scale.

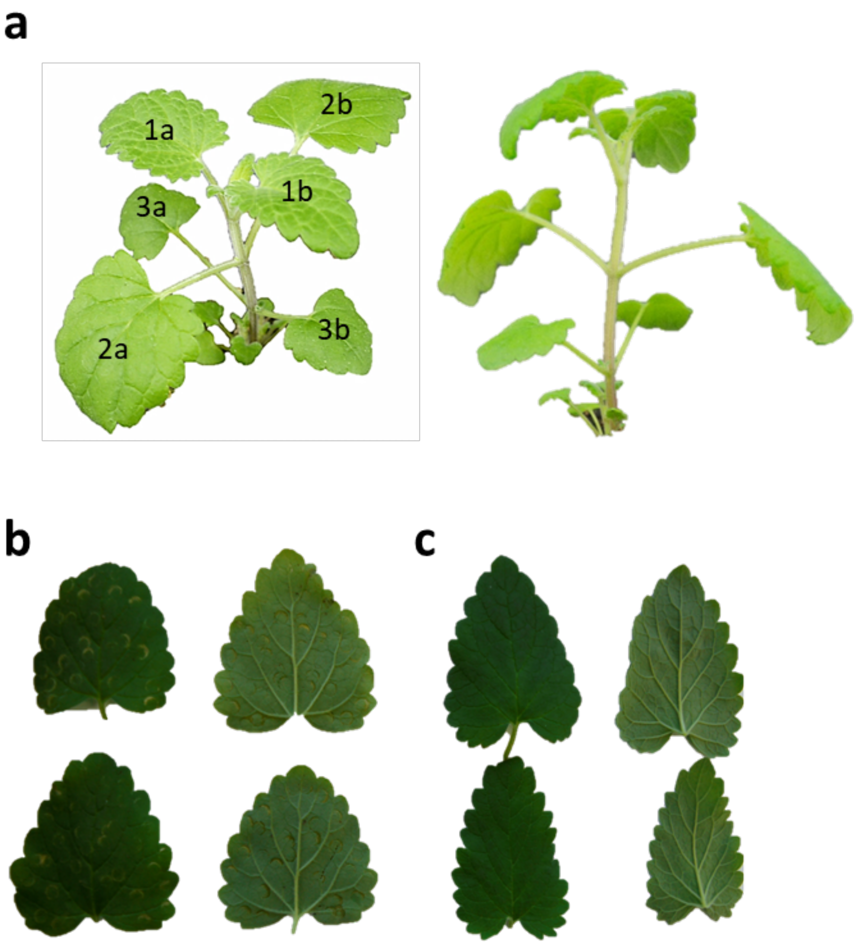


**Supplementary Fig. 3 (a)** Infiltration scheme of the upper six leaves (1a, 1b, 2a, 2b, 3a, 3b) of four-week-old *N. cataria* plants. **(b)** The infiltrated leaves showed very little necrosis at seven dpi compared to non-infiltrated leaves **(c)**.

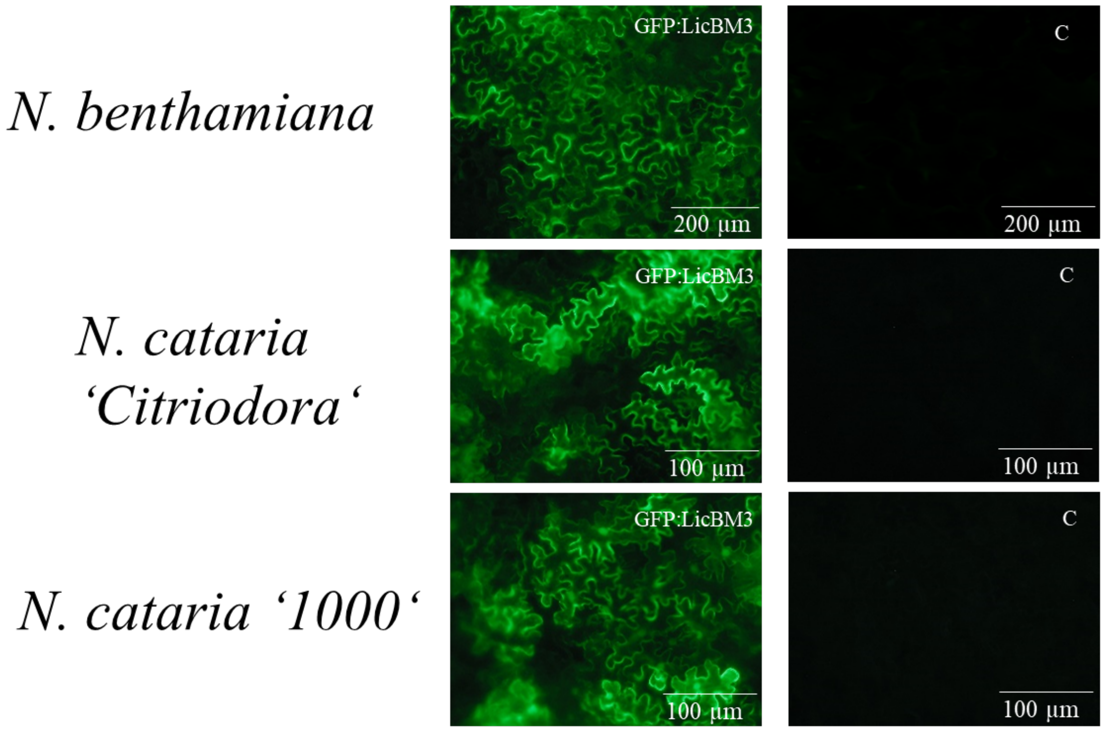


**Supplementary Fig. 4** Fluorescence microscopy of *N. cataria* variety 'Citriodora' and '1000' and *N. benthamiana* plants expressing *gfp:licBM3+P19*. Infiltration of plants with untransformed *Agrobacterium* served as a negative control (C).

### Additional information

#### Cloning of genes and generation of *A. tumefaciens* EHA105 and GV3101 for transient transformation of *N. cataria* and *N. benthamiana* plants

For introduction of relevant genes into the GoldenBraid (GB) system, GB-modified genes encoding for acyl–activating enzyme 1 (AAE1), olivetol synthase (OLS) and olivetolic acid cyclase (OAC) originating from *C. sativa* were synthesized by Integrated DNA Technologies (Coralville, IA, USA). The GoldenBraid assembly was then performed as described by Sarrion-Perdigones *et al.* (2013). Afterwards, the reaction mixtures were transformed into chemically competent *E. coli* TOP10 cells and positive clones were selected on solid medium. Subsequently, bacteria were grown in liquid LB, supplemented with appropriate antibiotics and plasmid DNA was extracted by means of the Plasmid Miniprep Kit I (VWR International GmbH, Darmstadt, Germany). All assemblies were verified by restriction analysis. The newly domesticated parts were sequenced by Eurofins Genomics using M13 primers: M13 uni (-21), 5'-TGT AAA ACG ACG GCC AGT-3' and M13 rev (-29), 5'-CAG GAA ACA GCT ATG ACC-3'. Finally, the desired Ω-level GB constructs were transformed into chemically competent *A. tumefaciens* EHA105 or GV3101 cells and the transfer confirmed by colony PCR using primer pairs (p35S-Cf3, 5'-CCA CGT CTT CAA AGC AAG TGG-3' and p35S-Cr4, 5'-TCC TCT CCA AAT GAA ATG AAC TTC C-3') screening for the cauliflower mosaic virus 35S promoter (P35S).

#### Fluorescence microscopy

Production of GFP-fused LicBM3 in *N. cataria* and *N. benthamiana* plants was analyzed using the Axioskop 40 Microscope (Zeiss, Oberkochen, Germany).

**Preparation of volatiles and GC–MS parameters**

For the analysis of metabolites in *N. cataria* plant material, the compounds were extracted with dichloromethane (DCM, Carl Roth, Karlsruhe, Germany) and the obtained samples measured with the QP 2010 Ultra gas-chromatography mass-spectrometry system (Shimadzu, Duisburg, Germany). Aliquots of 100 mg of frozen, powdered material were extracted with 2 ml of DCM. The extracts were shortly vortexed and sonicated for 10 min; filtration through a small glass column containing anhydrous $Na_2SO_4$ followed. The eluates were evaporated under a flow of nitrogen and resuspended in 100 µl of n-hexane (Carl Roth, Karlsruhe, Germany). Subsequently, 1 µl of each extract was injected onto the analytical GC–MS column (DB-5MS, 30 m 0.25 mm ID, 0.25 µm film thickness; Phenomenex, Aschaffenburg, Germany). Hydrogen was used as the carrier gas with a flow rate of 3.07 ml $min^{-1}$. Initial oven temperature was 40 °C (1 min holding); then, a linear gradient to 170 °C, at the rate of 7.5 °C $min^{-1}$, was applied and followed by a linear gradient to 300 °C at 60 °C $min^{-1}$, with the final temperature held for 5 min (total running time, 25.5 min). Ion source temperature was kept at 230 °C and the mass spectra of sample compounds were recorded by electron ionization at 70 eV, from 40 to 351 m/z.
